# Depression Attenuates Caudate and Dorsolateral Prefrontal Cortex Alpha and Beta Power Response to Reward

**DOI:** 10.1101/2024.02.05.578848

**Authors:** H Qian, GW Johnson, NC Hughes, DL Paulo, Z Zhao, D Subramanian, K Dhima, SK Bick

## Abstract

Depression is a prevalent psychiatric condition and a common comorbidity across neurological disorders. Common symptoms include anhedonia, negative emotional biases, and cognitive dysfunction. Beta (15-30 Hz) neural oscillations have been shown to increase during reward-based learning within fronto-striatal reward networks. Corticostriatal beta oscillations have also been implicated in cognitive functions including working memory. However, the relationship between beta oscillations and depression remains unknown. Using intracranial recordings, we aimed to investigate how depression modulates the spectral power of neural oscillations in corticostriatal structures during reward feedback in a working memory task.

Thirty movement disorder patients undergoing awake deep brain stimulation surgery with electrode trajectories traversing the caudate or dorsolateral prefrontal cortex (DLPFC) participated in this study. We recorded local field potential data intraoperatively as subjects completed a 2-back verbal working memory task where they identified whether a word matched the word presented two trials prior. Subjects received reward in the form of visual feedback for correct answers. Word stimuli had either a positive, negative, or neutral emotional valence. Subjects completed the Beck Depression Inventory-II preoperatively, and we used a cut-off score of 14 to identify patients with depression.

We found that caudate and DLPFC power increased in the alpha (8-15 Hz) and beta range during reward feedback and that this increase was significantly greater for subjects without depression compared to depressed subjects. In non-depressed patients, positive feedback stimuli evoked significantly higher beta power in the caudate during reward compared to neutral and negative stimuli. In depressed patients, emotional valence did not affect reward-related caudate spectral power, while DLPFC alpha power was significantly higher following positive emotional stimuli in comparison to neutral but not negative stimuli. We additionally found that anti-depressant medications (ADMs) generally blunted alpha and beta reward signaling processes in the DLPFC. This blunting effect on reward-related alpha power in the DLPFC, however, was reversed in depressed patients, indicating that the effects of ADMs on reward signaling processes may depend on whether a patient is exhibiting depression symptoms.

Our findings suggest that depression suppresses the alpha and beta power response to both reward and emotional stimuli during working memory, indicating power attenuation in these frequency bands may contribute to emotional and cognitive depression symptoms.

## Introduction

Depression is one of the most prevalent psychiatric conditions in the general population,^1^ and is a common and debilitating comorbidity in many neurological disorders including Parkinson’s disease, Alzheimer’s disease, multiple sclerosis, and stroke.^2,3^ Unfortunately, up to 30% of patients with depression are resistant to standard medical treatments.^4^ A better understanding of neural activity changes underlying depression is needed in order to develop improved treatment options for this disorder.

Anhedonia is a core feature of depression^5^ and is characterized by reward hyposensitivity and the inability to experience pleasure.^6,7^ Imaging research has shown that individuals with depression have altered reward processing, which may be associated with anhedonia. Dopaminergic neurons in the midbrain ventral tegmental area play an important role in reward signaling.^8^ These neurons project to multiple brain regions including caudate, nucleus accumbens, and prefrontal cortex, and as such information about reward is propagated to regions involved in learning, memory, and action selection.^9^ In healthy individuals, reward-related activity is seen with fMRI in multiple corticostriatal brain regions including nucleus accumbens, caudate, anterior cingulate, orbitofrontal cortex, and dorsolateral prefrontal cortex (DLPFC).^10–12^ Patients with major depressive disorder (MDD) have reduced activity and functional connectivity in frontal and striatal regions during reward anticipation.^7,13^ Several studies using fMRI have also found that subjects with MDD have attenuated neural activation in response to reward in multiple cortical and subcortical regions.^7,14–19^ Altered reward signaling in depression may be a biomarker of anhedonia. Decreased activity in frontal and striatal regions during reward anticipation has been correlated with anhedonia.^7^ This symptom is also associated with smaller fMRI reward signals in the caudate, nucleus accumbens, medial prefrontal cortex, and cingulate as well as reduced functional connectivity throughout fronto-striatal circuitry during reward delivery.^13,15,17,20^

Dysfunctional reward signaling may also contribute to cognitive symptoms of depression. Depression is thought to affect multiple domains of cognitive functioning including executive functioning, working memory, reinforcement learning, and emotional processing.^4,7,13,21–24^ Depressed patients have smaller striatal reward signals associated with impaired reinforcement learning.^20,25^ Smaller fMRI signals in the dorsomedial prefrontal cortex and striatum in response to reward are associated with deficits in executive control in MDD.^26^ Previous research with human subjects has shown that neural oscillations involved in working memory are significantly altered on a widespread scale in depressed individuals, particularly during working memory encoding, maintenance, and in response to increasing working memory load.^21–24^ However, the existing literature presents conflicting findings on whether depression alters working memory task performance,^21,23,27^ and it remains unclear if observed oscillatory changes result from depression-related cognitive deficits or instead reflect mechanisms meant to compensate for the cognitive effects of depression.

Emotion processing is also commonly altered in depression, with many depressed patients experiencing negative affective biases when processing emotional information.^28–30^ Patients with MDD have an impaired ability to recognize and remember positive valence emotional stimuli.^31^ Deficits in recognizing and processing emotional valence have been suggested to be biomarkers of anhedonia and may be associated with impairment in cognitive processes such as executive functioning and working memory.^31–34^ While healthy subjects have better working memory performance for positive valence emotional stimuli,^35^ subjects with MDD selectively attend to negative valence stimuli and have greater difficulty encoding positive stimuli.^33,34^ There have been mixed findings on how depression alters neural activation during emotion processing, but depressed individuals have generally been found to exhibit enhanced activation of emotion-sensitive prefrontal networks when processing negative information, including the ventromedial prefrontal cortex, ventral anterior cingulate cortex, and extended limbic system.^28,30^ Patients with depression also exhibit reduced activity in cognitive control and emotion regulation networks, including the dorsal anterior cingulate cortex, dorsomedial prefrontal cortex, and lateral prefrontal regions, during emotion processing tasks.^28–30,36^

A limitation of these prior bodies of work is that they have generally relied on functional imaging techniques, which have limited spatial and temporal resolution as they measure blood oxygenation changes rather than neural activity. These limitations can be overcome by intracranial EEG, which directly records local field potentials (LFPs) from brain structures with high temporal resolution. Previous work has shown that beta band neural oscillations in the caudate and DLPFC increase during reward-related feedback of cognitive tasks in human subjects.^37–39^ However, whether these corticostriatal reward-related neural oscillations are altered in depression is not known. Moreover, how reward and emotional processing may interact is not well understood. Finally, the impact of antidepressant medications, which modulate neurotransmitters involved in reward signaling, on the neural correlates of reward signaling is poorly understood. In this study, we used an emotional 2-back working memory task and intracranial recordings in human subjects undergoing deep brain stimulation surgery for movement disorders to investigate how reward-related caudate and DLPFC beta power increase may be modulated by depression and be related to anhedonia. We also investigated how emotional valence and antidepressant medications modulate reward signaling and may be modulated by depression.

## Methods and Materials

### Subjects

Thirty movement disorder patients participated in this study from 2021 to 2023. Patients underwent bilateral DBS surgery of the subthalamic nucleus (STN), globus pallidus internus (GPI), or ventral intermediate nucleus of the thalamus (VIM) under local anesthesia at our institution. All patients undergoing new DBS implantation with previously planned electrode trajectories contacting the caudate and/or DLPFC were eligible to participate in this study. Fourteen subjects had Parkinson’s disease, and 16 subjects had essential tremor. Demographic and clinical data for all subjects are detailed in **Table 1**. This study was approved by the Vanderbilt University Medical Center Institutional Review Board prior to initiation and all subjects provided written informed consent. We collected demographic, behavioral, and clinical data for all subjects from their electronic medical records.

**Table 1.** Patient demographic and disease-related information.

| Subject | Disorder | Target | Age (Years) | Sex | Handedness | Disease Duration (Years) | BDI-II | Antidepressants | UPDRS Score - Off | UPDRS Score - On | % Trials Correct | Caudate Channels | DLPCF Channels |
| --- | --- | --- | --- | --- | --- | --- | --- | --- | --- | --- | --- | --- | --- |
| 1 | PD | STN, bilateral | 56 | M | Right | 5 | 21 | None | 49 | 30 | 79.17 | 2 | 3 |
| 2 | PD | STN, bilateral | 62 | M | Right | 9 | 4 | Paroxetine (SSRI) | 53 | 28 | 87.67 | 3 | 2 |
| 3 | PD | STN, bilateral | 60 | F | Right | 11 | 0 | None | 49 | 35 | 52.74 | 3 | 3 |
| 4 | ET | VIM, bilateral | 43 | F | Right | 10 | 2 | None | 18 | 18 | 77.4 | 2 | 2 |
| 5 | ET | VIM, bilateral | 75 | F | Right | N/A | 7 | None | 18 | 19 | 54.79 | 1 | 2 |
| 6 | ET | VIM, bilateral | 72 | M | Right | N/A | 10 | Mirtazapine (Atypical) | 22 | 21 | 52.05 | 0 | 2 |
| 7 | PD | STN, bilateral | 58 | M | Right | 11 | 19 | Duloxetine (SNRI) | 60 | 5 | 74.07 | 0 | 6 |
| 8 | PD | STN, bilateral | 62 | M | Left | 5 | 12 | None | 41 | 10 | 87.67 | 2 | 3 |
| 9 | ET | VIM, bilateral | 68 | M | Right | 4 | 7 | None | 8 | 20 | 88.36 | 4 | 0 |
| 10 | ET | VIM, bilateral | 65 | F | Right | 13 | 16 | Doxepin (TCA), Bupropion (Atypical) | N/A | N/A | 76.79 | 0 | 4 |
| 11 | PD | STN, bilateral | 56 | M | Right | 6 | 8 | None | 71 | N/A | 84.93 | 3 | 3 |
| 12 | ET | VIM, bilateral | 68 | F | Right | 38 | 2 | None | 11 | 18 | 71.23 | 0 | 2 |
| 13 | PD | STN, bilateral | 64 | M | Right | 8 | 4 | None | 30 | 2 | 72.6 | 2 | 3 |
| 14 | ET | VIM, bilateral | 71 | M | Left | 10 | 9 | None | 18 | 18 | 82.19 | 2 | 1 |
| 15 | PD | STN, bilateral | 60 | M | Right | 7 | 20 | None | 32 | 24 | 84.25 | 3 | 0 |
| 16 | PD | GPI, bilateral | 56 | M | Right | 15 | 7 | None | 47 | 25 | 84.93 | 0 | 6 |
| 17 | PD | GPI, bilateral | 57 | M | Right | 5 | 9 | None | 33 | 12 | 78.77 | 0 | 5 |
| 18 | PD | GPI, bilateral | 76 | M | Right | 9 | 18 | Sertraline (SSRI) | 60 | 45 | 49.32 | 2 | 3 |
| 19 | ET | VIM, bilateral | 73 | F | Right | 20 | 8 | Sertraline (SSRI) | 21 | 19 | 79.45 | 1 | 0 |
| 20 | PD | STN, bilateral | 67 | M | Right | 5 | 22 | None | 30 | 10 | 78.77 | 2 | 3 |
| 21 | ET | VIM, bilateral | 63 | M | Right | 5 | 7 | Sertraline (SSRI) | 18 | 25 | 73.97 | 0 | 4 |
| 22 | ET | VIM, bilateral | 55 | M | Right | 4 | 24 | None | 0 | 12 | 63.7 | 0 | 2 |
| 23 | ET | VIM, bilateral | 66 | M | Right | 9 | 10 | Duloxetine (SNRI) | 17 | 13 | 78.77 | 0 | 4 |
| 24 | PD | GPI, bilateral | 77 | F | Right | 12 | 3 | None | 53 | 23 | 65.75 | 0 | 6 |
| 25 | PD | GPI, bilateral | 69 | M | Right | 10 | 9 | None | 28 | 15 | 79.17 | 1 | 3 |
| 26 | ET | VIM, bilateral | 52 | M | Right | 31 | 1 | None | 17 | 7 | 70.83 | 1 | 2 |
| 27 | ET | VIM, bilateral | 76 | M | Right | 21 | 5 | None | 22 | 16 | 68.75 | 2 | 2 |
| 28 | ET | VIM, bilateral | 71 | F | Right | 15 | 2 | Citalopram (SSRI) | 11 | 11 | 77.08 | 2 | 0 |
| 29 | ET | VIM, bilateral | 63 | M | Right | N/A | 0 | None | 17 | 15 | 87.5 | 1 | 2 |
| 30 | ET | VIM, bilateral | 44 | M | Right | 4 | 9 | None | 21 | 18 | 52.08 | 1 | 1 |
| Total / Average ± SD | 14 PD, 16 ET | 9 STN, 5 GPI, 16 VIM | 63.5 ± 8.85 | 22 M, 8 F | 28 Right, 2 Left | 11.19 ± 8.18 | 7 D, 23 ND | 9 ADM, 21 no ADM | 30.17 ± 18.09 | 18.36 ± 9.18 | 73.83 ± 11.69 | 40 | 79 |

### Preoperative Evaluation

Preoperatively, subjects completed the Beck Depression Inventory-II (BDI-II) as a self-reported measure of their depressive symptoms. We categorized subjects as depressed if their total BDI-II score was 14 or higher.^40,41^ In order to evaluate specific depression symptoms, we also computed affective (encompassing mood), somatic (encompassing bodily and cognitive difficulties), and anhedonia (encompassing loss of interest and pleasure) symptom subscale scores from BDI-II items as has previously been described.^42–45^ Items included in each factor are listed in **Table 2**.

**Table 2.** BDI-II factor structure and subdomain items.

| Item Number | Item Description | Anhedonia | Affective | Somatic |
| --- | --- | --- | --- | --- |
| 1 | Sadness |  | ✓ |  |
| 2 | Pessimism |  | ✓ |  |
| 3 | Past Failure |  | ✓ |  |
| 4 | Loss of Pleasure | ✓ | ✓ |  |
| 5 | Guilty Feelings |  | ✓ |  |
| 6 | Punishment Feelings |  | ✓ |  |
| 7 | Self-Dislike |  | ✓ |  |
| 8 | Self-Criticalness |  | ✓ |  |
| 9 | Suicidal Thoughts or Wishes |  | ✓ |  |
| 10 | Crying |  |  | ✓ |
| 11 | Agitation |  | ✓ |  |
| 12 | Loss of Interest | ✓ | ✓ |  |
| 13 | Indecisiveness |  |  | ✓ |
| 14 | Worthlessness |  | ✓ |  |
| 15 | Loss of Energy |  |  | ✓ |
| 16 | Change in Sleep Pattern |  |  | ✓ |
| 17 | Irritability |  | ✓ |  |
| 18 | Changes in Appetite |  |  | ✓ |
| 19 | Concentration Difficulty |  |  | ✓ |
| 20 | Tiredness of Fatigue |  |  | ✓ |
| 21 | Loss of Interest in Sex | ✓ |  | ✓ |

### Surgery

Patients underwent bilateral DBS electrode implantation surgery under local anesthesia with clinical microelectrode recordings. Dopaminergic medications for Parkinson’s disease patients were held the night prior to surgery according to standard clinical protocol to facilitate intraoperative motor testing. Per our standard clinical protocol, a custom-made mini-stereotactic frame (FHC Inc., Bowdoin, ME) was mounted with two microdrives. Each microdrive held two or three microelectrodes depending on the final DBS target. Macro contacts on the microelectrode protective tube 10 mm from the micro tip allowed recording of local field potentials (LFPs) (FHC Inc., Bowdoin, ME). This setup allowed for simultaneous bilateral recordings. LFPs were recorded via the FHC Guideline 5 system with a sampling rate of 1000 Hz. Twenty subjects had recording electrodes that traversed the caudate, and 26 subjects had electrodes that traversed the DLPFC for a total of 40 caudate and 79 DLPFC individual channel recordings.

### Cognitive Task

At the beginning of the research session, two minutes of LFP data was recorded while subjects rested quietly with their eyes closed. Subjects then participated in an emotional verbal 2-back working memory task, during which they were sequentially visually presented with a series of words characterized for their emotional qualities **(Fig. 1A)**. For each word, a response cue appeared following a 1000 ms pause prompting the subject to respond whether the word presented during the current trial matched the word presented two trials prior by pressing a button. The side of the button assigned to yes versus no response was randomly assigned on a trial-by-trial basis and indicated on the screen. Following response, subjects received visual feedback for 500 ms, where the word turned green to indicate a correct response or red to indicate an incorrect response. Each trial had a 50% chance of being a “match,” in which the word presented matched the word presented two trials prior.

**Figure 1.**
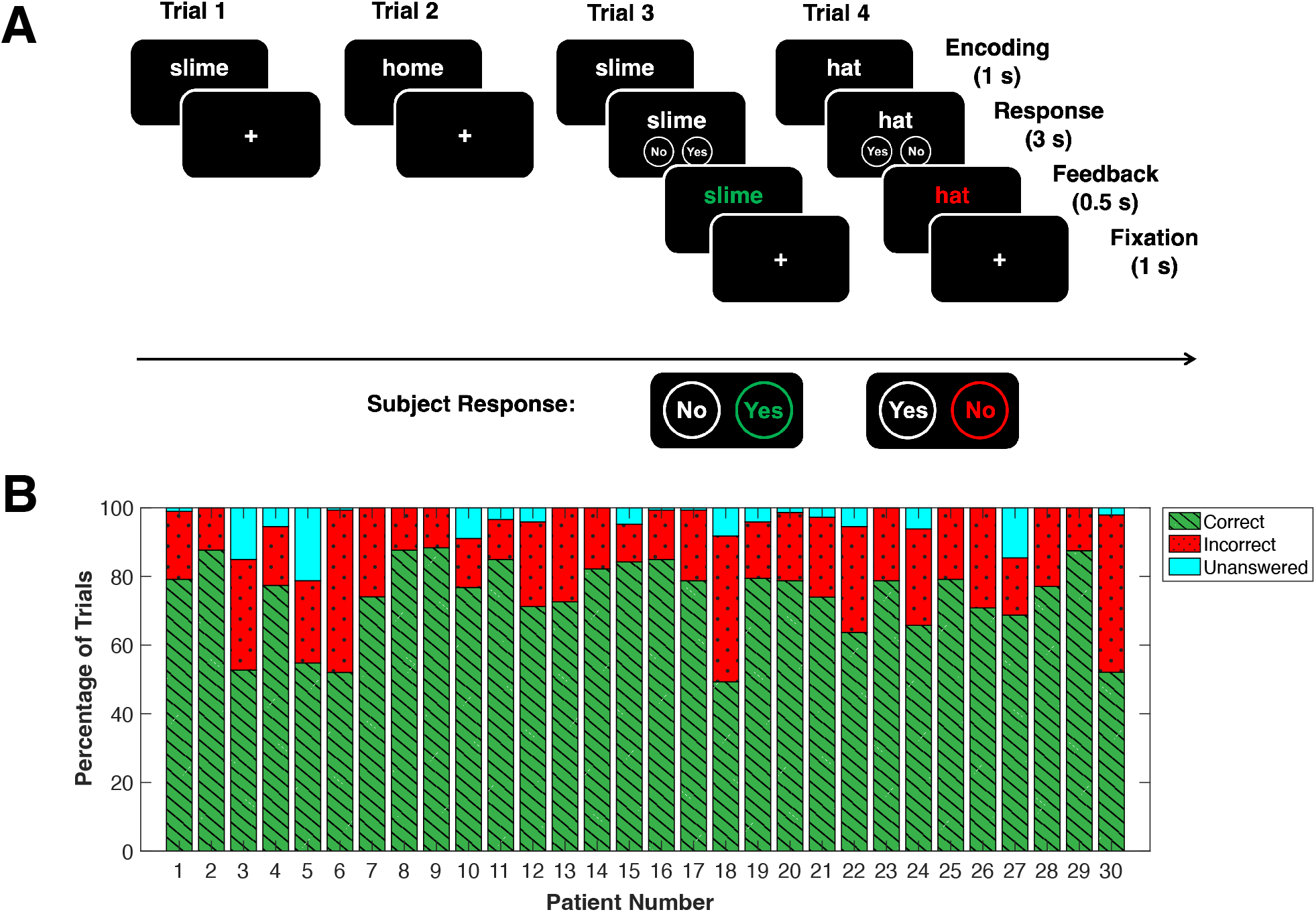
2-back working memory task and task performance. **(A)** 2-back verbal working Memory Task: Word stimuli is presented to the subject for 1 second, followed by a response cue prompting the subject to answer whether the current word matches the word presented two trials prior. The orientation of the buttons corresponding to Yes/No is randomly assigned on a trial by trial basis. Following the response, visual feedback is presented for 0.5 s. Green represents correct response, red represents incorrect response. **(B)** Task performance by subject: Percentage of correct (green), incorrect (red), and unanswered (cyan) trials per subject, averaged across blocks.

In order to examine emotion-related LFP and working memory characteristics, we used words selected from the Affective Norms for English Words (ANEW) dataset of words with previously characterized emotional features^46^ **(Table 3)**. In this dataset, words are rated on a scale from 1-9 for valence (negative to positive characteristics), arousal (degree of feeling calm to excited), and dominance (degree of feeling controlled to feeling in control). For each block of the task, there was a predetermined set of four negative words, four neutral words, and four positive words. There were three different blocks with 12 unique emotional words each. We selected positive and negative valence words to have similar degrees of deviation from neutral valence (5 by definition) and similar arousal ratings. Mean valence was 2.56 ± 0.33 for negative valence words, 5.46 ± 0.30 for neutral valence words, and 7.79 ± 0.36 for positive valence words. As expected, valence significantly differed between words across the three emotions upon analysis with a one-way ANOVA [*F* = 755, *d.f.* = 2, *P* = 2.79E-28]. By a Wilcoxon rank-sum comparison, deviation in valence from neutral was slightly greater for positive words compared to negative words [*P* = 0.021, *z* = -2.3]. Dominance also significantly differed between all three emotion categories [*F* = 31, *d.f.* = 2, *P* = 3.1E-08], with mean dominance being 4.48 ± 0.30 for negative words, 5.10 ± 0.33 for neutral words, and 5.69 ± 0.49 for positive words. Mean arousal was 5.36 ± 0.69 for negative words, 4.67 ± 0.64 for neutral words, and 5.23 ± 0.61 for positive words, which differed between negative and neutral words [*F* = 3.8, *d.f.* = 2, *P* = 0.032]. Subjects completed two 75-trial blocks of this task.

**Table 3.**
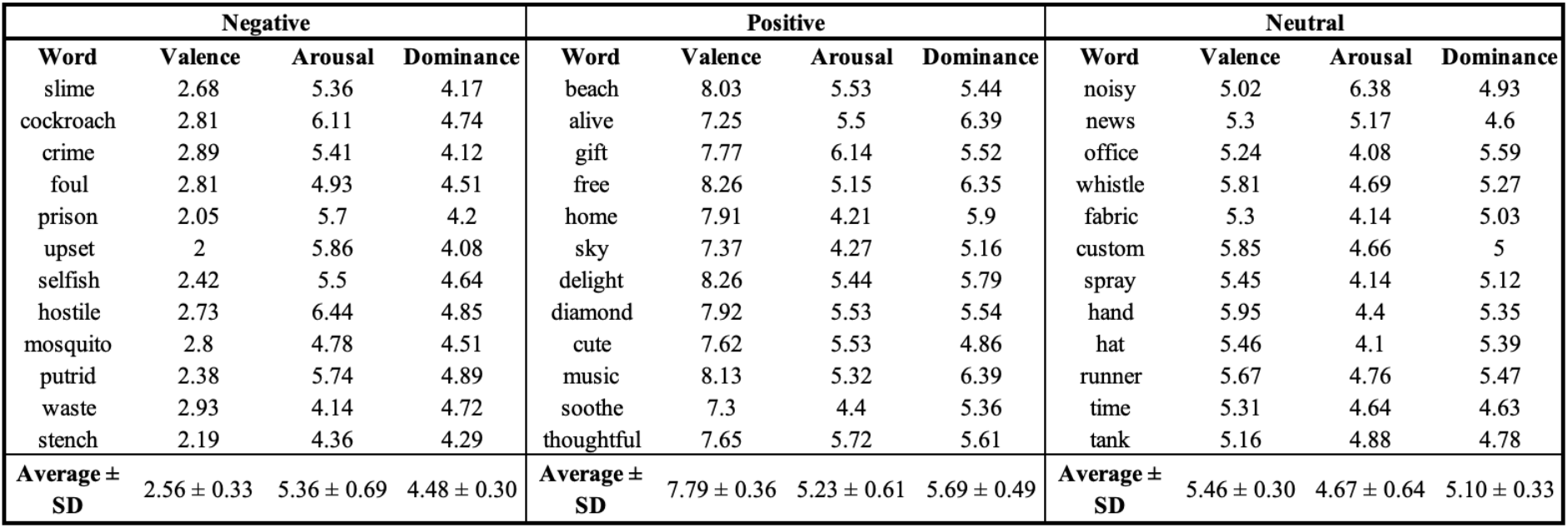
Emotional valence of task word stimuli.

| Negative |  |  |  | Positive |  |  |  | Neutral |  |  |  |
| --- | --- | --- | --- | --- | --- | --- | --- | --- | --- | --- | --- |
| Word | Valence | Arousal | Dominance | Word | Valence | Arousal | Dominance | Word | Valence | Arousal | Dominance |
| slime | 2.68 | 5.36 | 4.17 | beach | 8.03 | 5.53 | 5.44 | noisy | 5.02 | 6.38 | 4.93 |
| cockroach | 2.81 | 6.11 | 4.74 | alive | 7.25 | 5.5 | 6.39 | news | 5.3 | 5.17 | 4.6 |
| crime | 2.89 | 5.41 | 4.12 | gift | 7.77 | 6.14 | 5.52 | office | 5.24 | 4.08 | 5.59 |
| foul | 2.81 | 4.93 | 4.51 | free | 8.26 | 5.15 | 6.35 | whistle | 5.81 | 4.69 | 5.27 |
| prison | 2.05 | 5.7 | 4.2 | home | 7.91 | 4.21 | 5.9 | fabric | 5.3 | 4.14 | 5.03 |
| upset | 2 | 5.86 | 4.08 | sky | 7.37 | 4.27 | 5.16 | custom | 5.85 | 4.66 | 5 |
| selfish | 2.42 | 5.5 | 4.64 | delight | 8.26 | 5.44 | 5.79 | spray | 5.45 | 4.14 | 5.12 |
| hostile | 2.73 | 6.44 | 4.85 | diamond | 7.92 | 5.53 | 5.54 | hand | 5.95 | 4.4 | 5.35 |
| mosquito | 2.8 | 4.78 | 4.51 | cute | 7.62 | 5.53 | 4.86 | hat | 5.46 | 4.1 | 5.39 |
| putrid | 2.38 | 5.74 | 4.89 | music | 8.13 | 5.32 | 6.39 | runner | 5.67 | 4.76 | 5.47 |
| waste | 2.93 | 4.14 | 4.72 | soothe | 7.3 | 4.4 | 5.36 | time | 5.31 | 4.64 | 4.63 |
| stench | 2.19 | 4.36 | 4.29 | thoughtful | 7.65 | 5.72 | 5.61 | tank | 5.16 | 4.88 | 4.78 |
| Average ± SD | 2.56 ± 0.33 | 5.36 ± 0.69 | 4.48 ± 0.30 | Average ± SD | 7.79 ± 0.36 | 5.23 ± 0.61 | 5.69 ± 0.49 | Average ± SD | 5.46 ± 0.30 | 4.67 ± 0.64 | 5.10 ± 0.33 |

### Neurophysiology and Statistical Analysis

LFP analysis was performed offline using MATLAB (MathWorks, Natick MA) and FieldTrip MATLAB toolbox.^47^ Recordings were visually examined for noise, and trials and/or channels were excluded if the signal was contaminated by significant artifact. Data was notch filtered at 60 Hz to remove line noise, high-pass filtered at 1 Hz, and aligned to task events using digital event triggers. Spectral power was calculated from 1-100 Hz using the Morlet wavelet time-frequency transformation in FieldTrip. For each block, task-based power was z-scored across all trials for each channel and frequency interval, while resting power was log-transformed by base 10 and averaged over time.

To test our primary hypothesis on the relationship between depression and reward-induced power in working memory, we analyzed the spectral power of neural oscillations in the caudate and DLPFC during the 1000 ms period following the onset of visual feedback. We first used cluster-based permutation testing to identify time-frequency clusters of spectral power that significantly differed between feedback for correct (rewarded) vs incorrect responses (non-rewarded).^48,49^ For each channel, we first averaged spectral power across rewarded and non-rewarded trials separately. Our cluster-based permutation analysis then used a paired t-test to generate a T-value to compare power for rewarded and non-rewarded trials at each time point and frequency interval from 1-100 Hz and then summed the paired T-values within clusters with adjacent data points above the statistical significance threshold of 0.05. The labels designating averaged channel-level recordings as corresponding to rewarded versus non-rewarded trials were then randomly reassigned 1000 times with the T-statistic similarly determined. Significant clusters were defined as those with summed T-values greater than that of 95% of the randomly determined comparisons. For each frequency band that fell within the frequency range of a significant frequency-by-time cluster, we then performed a one-dimensional cluster-based permutation test to identify time clusters in which band-averaged power differed between correct and incorrect trials. We used the time clusters identified in this one-dimensional cluster-based permutation analysis as our windows of interest to compare average spectral power in each frequency band across stimuli, trial types, and patient groups of interest. For all subsequent statistical analyses, we averaged band-specific power over these time windows.

To analyze the general effect of reward feedback on neural oscillatory power, we performed a Wilcoxon signed rank analysis on the time-averaged power of rewarded versus non-rewarded trials within frequency bands of interest for all subjects. We additionally classified the 500 ms before feedback presentation as a baseline period and conducted a Wilcoxon signed rank test to analyze changes in neural oscillatory power during reward feedback compared to baseline.

To investigate the group effects of depression status on neural oscillatory power, we conducted a Wilcoxon rank sum test to determine whether averaged power during feedback for correct trials significantly differed by patient group. Separating patients into depressed and non-depressed groups, we used a one-way repeated measures ANOVA to assess the effect of the emotional valence of feedback stimuli on average spectral power. We used a one-way repeated measures ANOVA to assess the effect of the emotional valence of stimuli on average spectral power during reward for depressed and non-depressed groups separately. For significant ANOVA results, we used the Tukey-Kramer post hoc test to evaluate the significance of pair-wise comparisons. Additionally, we used the Pearson correlation coefficient to assess the relationship between spectral power during reward feedback and each subject’s total BDI-II score, as well as their anhedonia, affective factor, and somatic factor subscores. We conducted correlation analyses on the subject level, averaging all channels per recording region for each subject, while all other statistical analysis on spectral power data was conducted on the channel level.

We analyzed subjects’ behavioral data in conjunction with their neurophysiological data to explore whether depression was associated with working memory task performance and reaction time. We standardized reaction time comparisons across subjects by z-scoring reaction times across all trials of a block. We used the Pearson correlation coefficient to assess how each subject’s percentage of correct trials was associated with BDI-II score as well as caudate and/or DLPFC alpha and beta power following feedback for rewarded trials. We also used the Wilcoxon rank sum test to assess differences in task performance between depressed and non-depressed patients. The same statistical analyses were conducted to investigate the effects of depression on reaction time. We additionally performed a two-way repeated measures ANOVA to examine the interaction between depression and reaction time for correct and incorrect trials.

As an exploratory analysis, we were also interested in how antidepressant medications (ADMs) may impact reward-related oscillatory power. We categorized patients as taking ADMs if they were routinely taking serotonin reuptake inhibitors (SSRIs), serotonin–norepinephrine reuptake inhibitors (SNRIs), tricyclic antidepressants (TCAs), monoamine oxidase inhibitors (MAOIs), or atypical antidepressants at the time of surgery. Medications that met this inclusion criteria are detailed in **Table 1**. We evaluated the effects of antidepressant medication using a rank sum test to compare average spectral power following reward feedback for patients who were taking ADMs and those not taking ADMs. We also performed a two-way ANOVA with antidepressant medication status and depression status as between-subjects factors to determine whether there was a significant interaction between these demographic characteristics.

All P-values were Bonferroni-corrected for multiple comparisons based on the number of frequency bands and/or behavioral measures assessed.

## Results

### Behavioral Data

All subjects completed both resting data collection and the 2-back emotional verbal working memory task. Average working memory task performance was 73.8% correct, 22.5% incorrect, and 3.70% unanswered trials **(Fig. 1B)**. There was no relationship between BDI-II score and task performance on the 2-back task [*R* = -0.019, *P* = 0.92], nor did we find a significant difference in task performance as determined by mean percent correct trials between depressed and non-depressed groups [*P* = 0.52, *z* = -0.64]. Task performance on trials in which subjects responded to positive [*P* = 0.52, *z* = -0.64], neutral [*P* = 0.49, *z* = -0.69], and negative [*P* = 0.77, *z* = 0.29] word stimuli also did not significantly differ based on depression status.

Average z-scored reaction time during incorrect trials was significantly faster for depressed patients compared to non-depressed patients [*P* = 0.027, *z* = -2.9]. We found a significant interaction between depression and trial accuracy (correct versus incorrect) on reaction time [*F* = 4.2, *d.f.* = 1, *P* = 0.049], with non-depressed patients responding significantly slower on incorrect trials compared to correct trials [*P* = 0.0013]. There was no significant pairwise difference between reaction time for correct and incorrect trials for depressed patients [*P* = 0.71].

### Caudate and DLPFC Neural Oscillations During Reward

Following visual feedback for correct trials, both the caudate and DLPFC exhibited a significant increase in oscillatory power in the low to mid-frequency range. This increase in power was not observed following feedback for incorrect trials. In the caudate, the frequency-by-time cluster in which the spectral power of rewarded and non-rewarded trials significantly differed ranged from 8-44 Hz and 280-1000 ms after feedback appearance [*P* = 0.0010]. In the DLPFC, this frequency-by-time cluster ranged from 7-33 Hz and 20-1000 ms after feedback appearance [*P* = 0.0010]. We also identified an additional frequency-by-time cluster where rewarded versus non-rewarded spectral power significantly differed in the DLPFC ranging from 36-100 Hz and 240-570 ms after feedback appearance [*P* = 0.0010]. Although this cluster met our threshold for significance, we chose to focus our subsequent analysis on the frequency-by-time cluster with the greatest significant difference **(Fig. 2A, 2D)**.

**Figure 2.**
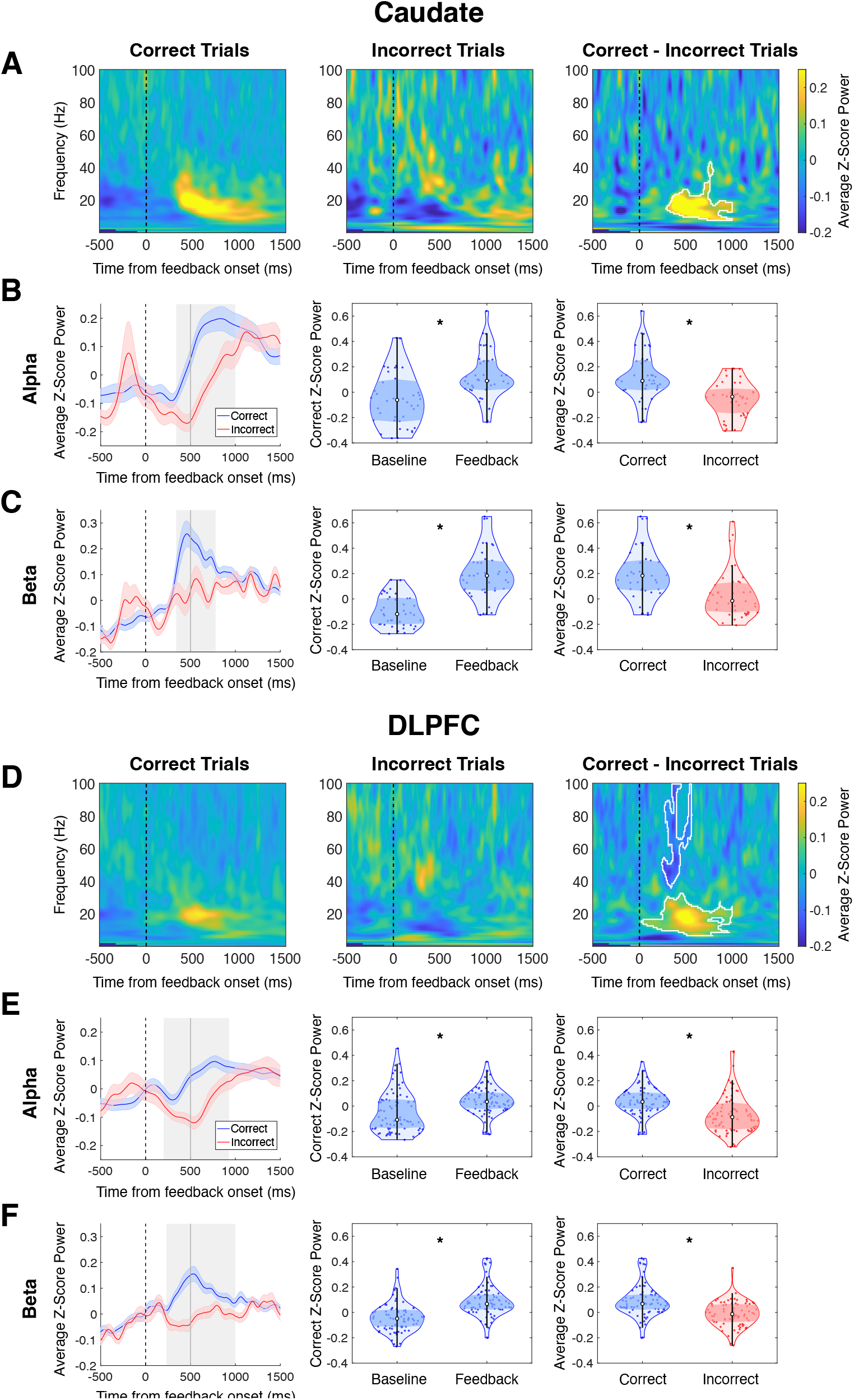
Caudate and DLPFC spectral power during reward feedback. Caudate and DLPFC data is shown in the first three rows (A-C) and the last three rows (D-F) respectively. **A and D:** Spectrograms of z-scored power averaged over correct (left) trials, incorrect (center), and the difference between average z-scored power for correct and incorrect trials (right). Dotted line indicates presentation of reward feedback. Significantly different time-frequency clusters (p < 0.05) between correct and incorrect trials are outlined in white. **B, C, E, F: (Left)** Time courses of average z-scored alpha power (B, E) and beta power (C, F) during feedback for correct and incorrect trials. Shading around the average line indicates standard error. Dotted line indicates feedback onset and solid line indicates end of feedback presentation. Gray shaded boxes indicate time clusters where band-specific power was significantly different between correct and incorrect power. **(Center)** Violin plots comparing average correct z-scored alpha (B, E) and beta (C, F) power at baseline (500 ms before feedback onset) and during time clusters of significant difference between correct and incorrect trials. White circle indicates median, middle shaded region indicates interquartile range, and black line indicates the data minimum and maximum excluding outliers. Beta and alpha power in both caudate and DLPFC significantly increased following feedback for correct trials. **(Right)** Violin plots comparing average z-scored alpha (B, E) and beta (C, F) power for correct and incorrect trials during significant time clusters. Beta and alpha power in both caudate and DLPFC was significantly higher during feedback for correct trials compared to incorrect trials. * = p <0.05. DLPFC = dorsolateral prefrontal cortex.

For both the caudate and DLPFC, the most significant frequency-by-time clusters identifying rewarded versus non-rewarded trial differences encompassed only the alpha (8-15 Hz) and beta (15-30 Hz) frequency bands in their entirety. We therefore performed additional analysis to determine the time range in which oscillatory power within each of these frequency bands was significantly different between rewarded and non-rewarded trials. In the caudate, averaged alpha band spectral power significantly differed between rewarded and non-rewarded trials from 340-1000 ms after feedback appearance [*P =* 0.0020] **(Fig. 2B)**. In the beta band, this differed from 340-780 ms following feedback appearance [*P =* 0.0020] **(Fig. 2C)**. In the DLPFC, spectral power was significantly different between rewarded and non-rewarded trials from 200-930 ms following feedback appearance in the alpha band [*P =* 0.0020] **(Fig. 2E)** and from 230-1000 ms following feedback appearance in the beta band [*P =* 0.0020] **(Fig. 2F)**.

We evaluated whether the power differences between rewarded and non-rewarded trials during feedback were associated with task-related activity by comparing oscillatory power within the time windows of interest to baseline power. In both caudate and DLPFC, oscillatory power for both alpha [*P* = 6.3E-04, *z* = -3.6 caudate and *P* = 9.8E-06, *z* = -4.6 DLPFC] and beta [*P* = 3.7E-07, *z* = -5.2 caudate and *P* = 4.0E-10, *z* = -6.4 DLPFC] bands significantly increased from baseline following feedback for rewarded trials. Beta power increased to a lesser degree from baseline following feedback for non-rewarded trials [*P* = 2.5E-02, *z* = -2.5 caudate and *P* = 9.5E-03, *z* = -2.8 DLPFC], while alpha power did not significantly differ from baseline [*P* = 1.4, *z* = -0.39 caudate and *P* = 3.2E-01, *z* = 1.4 DLPFC]. Average alpha power [*P* = 1.3E-05, *z* = 4.5 caudate and *P* = 1.4E-06, *z* = 5.0 DLPFC] and beta power [*P* = 9.7E-08, *z* = 5.5 caudate and *P* = 1.9E-08, *z* = 5.7 DLPFC] were also significantly higher following feedback for rewarded trials compared to non-rewarded trials, suggesting that reward significantly increases alpha and beta power in these structures **(Fig. 2B, 2C, 2E, 2F)**.

Task performance did not correlate with either alpha [*R* = 0.0070, *P* = 2.0 caudate and *R* = 0.0026, *P* = 1.98 DLPFC] or beta [*R* = 0.16, *P* = 0.98 caudate and *R* = 0.21, *P* = 0.60 DLPFC] power following reward presentation. Z-scored reaction times for correct trials also did not significantly correlate with alpha [*R* = -0.34, *P* = 0.30 caudate and *R* = -0.23, *P* = 0.50 DLPFC] or beta [*R* = -0.47, *P* = 0.072 caudate and *R* = -0.29, *P* = 0.32 DLPFC] power following reward.

### Depression and Reward-Related Oscillatory Power

Compared to non-depressed patients, depressed patients exhibited significantly lower beta power in both the caudate [*P* = 1.2E-04, *z* = -4.0] and DLPFC [*P* = 6.5E-05, *z* = -4.2] following reward **(Fig. 3B)**. Caudate alpha power was also significantly lower for depressed patients compared to non-depressed patients [*P* = 4.7E-03, *z* = -3.0]. Following non-reward feedback, we found no significant difference in alpha power [*P* = 1.5, *z* = 0.32 caudate and *P* = 2.0, *z* = -0.0056 DLPFC] or DLPFC beta power [*P* = 1.4E-01, *z* = -1.8] between patient groups **(Fig. 3B)**. However, caudate beta power following non-reward feedback was also significantly lower in depressed patients compared to non-depressed patients [*P* = 3.9E-02, *z* = -2.3].

**Figure 3.**
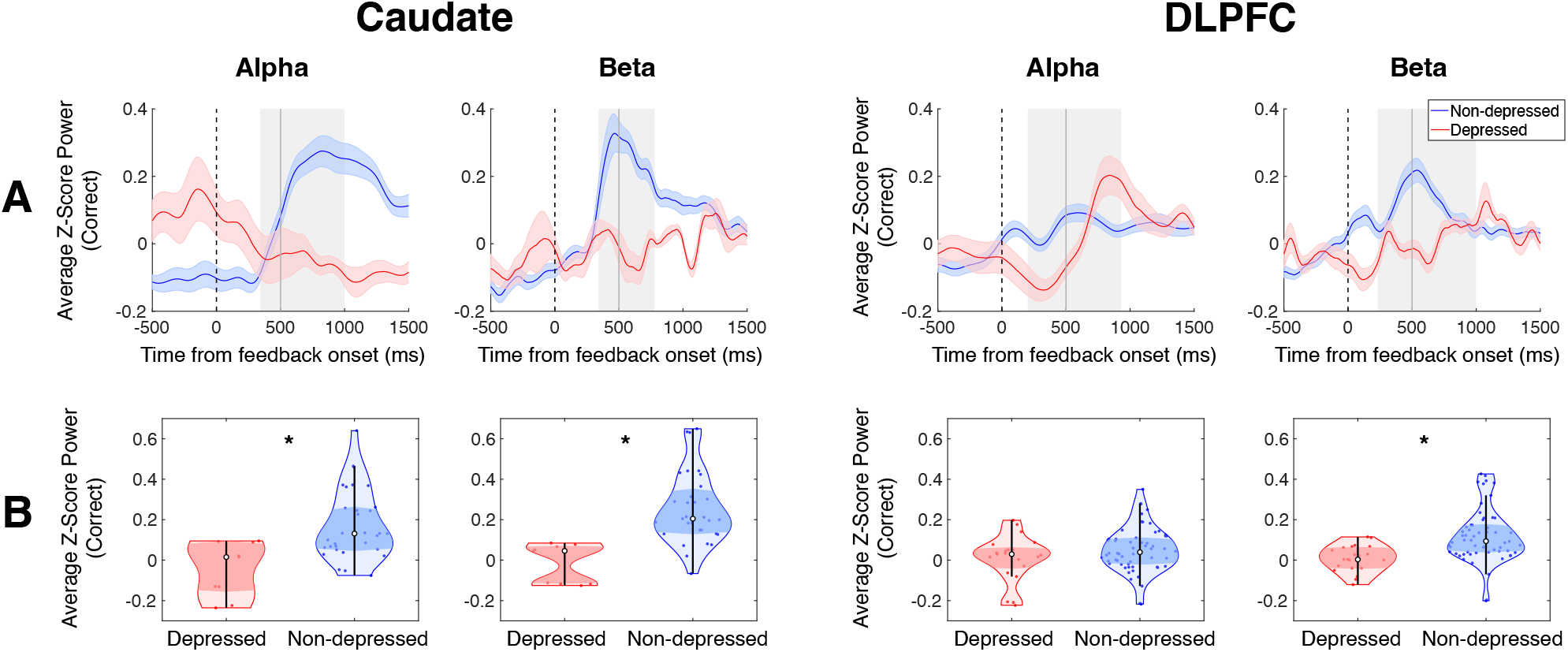
Effects of depression on caudate and DLPFC alpha and beta power during reward feedback. From left to right, caudate and DLPFC data is shown in the first two columns and the last two columns respectively. **(A)** Time courses of average z-scored alpha and beta power during reward feedback for correct trials for depressed and non-depressed patients. Shading around the average line indicates standard error. Dotted line indicates feedback onset and solid line indicates end of feedback presentation. Gray shaded boxes indicate time clusters where band-specific power was significantly different between correct and incorrect power across all subjects. **(B)** Violin plots comparing average correct z-scored beta power between depressed and non-depressed patients during time clusters of significant difference between correct and incorrect trials. White circle indicates median, middle shaded region indicates interquartile range, and black line indicates the data minimum and maximum excluding outliers. In both caudate and DLPFC, average beta power during reward feedback was significantly higher for non-depressed patients compared to depressed patients. Caudate alpha power was also significantly higher for non-depressed patients. * = p <0.05. DLPFC = dorsolateral prefrontal cortex.

To investigate the relationship between reward-related alpha and beta power increases and depression severity and symptoms, we examined the relationship between reward-related power and total BDI-II scores, as well as anhedonia, somatic, and affective subscores. Caudate beta power following reward was inversely correlated with total BDI-II score [*R* = -0.61, *P* = 0.014], anhedonia subscore [*R* = -0.55, *P* = 0.035], and somatic factor subscore [*R* = -0.62, *P* = 0.011] but was not associated with Affective factor subscore [*R* = -0.52, *P* = 0.059]. DLPFC beta power did not show a significant correlation with either total BDI-II score or its subscores. Alpha power in both DLPFC and caudate was also not correlated with BDI-II scores **(Fig. 4)**. These findings suggest that lower reward-related beta power in the caudate is associated with overall depression severity as well as greater anhedonia and somatic symptoms.

**Figure 4.**
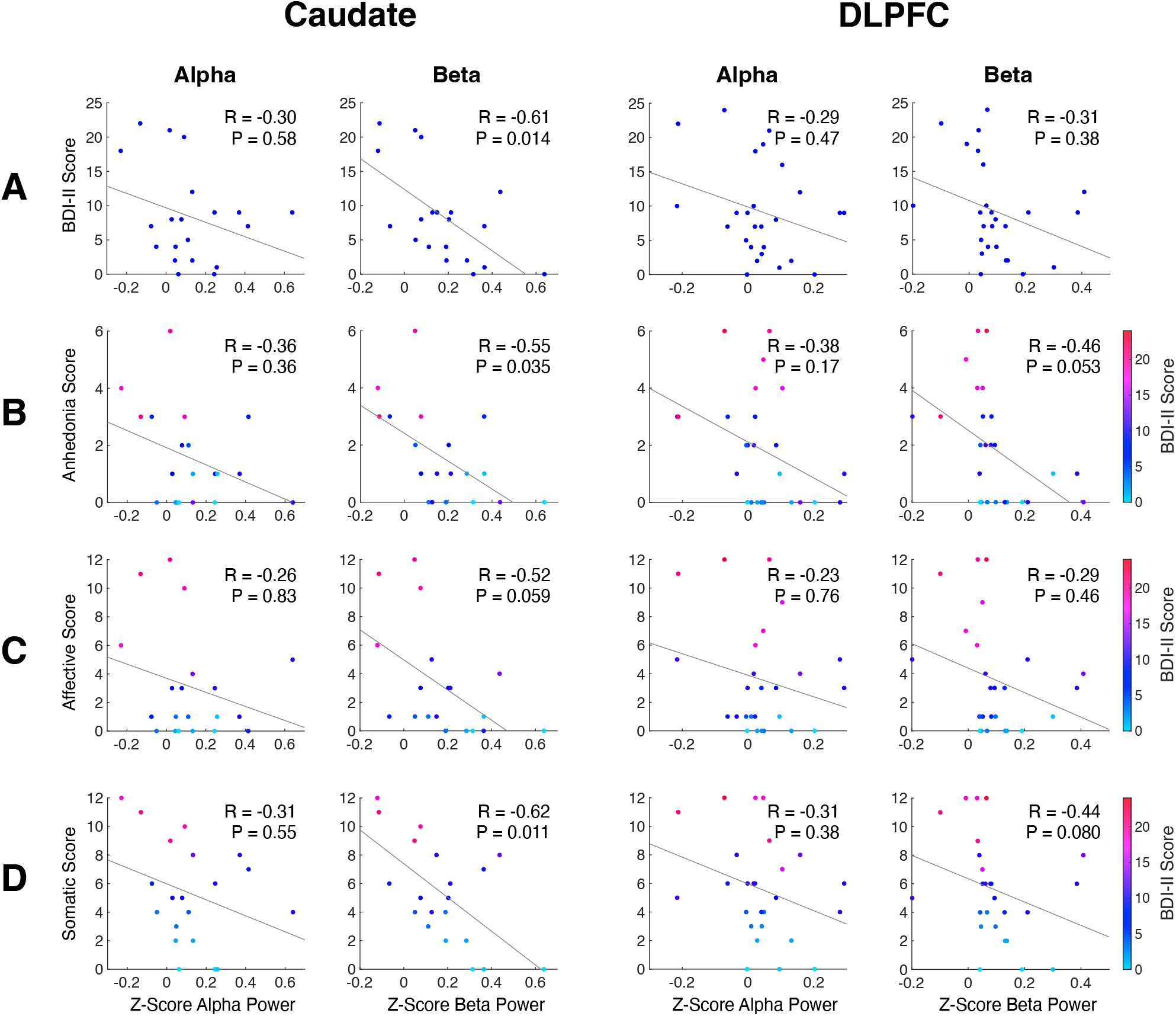
Correlations between BDI-II subdomain scores and caudate and DLPFC power during reward feedback. From left to right columns, scatterplots with correlations are shown with caudate alpha power, caudate beta power, DLPFC alpha power, and DLPFC beta power, respectively. **(A)** Correlations between total BDI-II score and average z-score power following feedback for rewarded trials. **(B)** Correlations between anhedonia subscore of BDI-II and average z-score power following feedback for rewarded trials. **(C)** Correlations between affective subscore of BDI-II and average z-score power following feedback for rewarded trials. **(D)** Correlations between somatic subscore of BDI-II and average z-score power following feedback for rewarded trials. Caudate beta power following reward exhibited a significant negative correlation with subjects’ total BDI-II score, anhedonia subscore, and somatic factor subscore.

From these results, we additionally analyzed caudate and DLPFC spectral power during resting-state recordings to determine whether depression also altered corticostriatal spectral power outside of the context of a cognitive task. Using the Wilcoxon rank sum comparison, we compared caudate and DLPFC resting power between depressed and non-depressed patients across the delta (1-4 Hz), theta (3-8 Hz), alpha, beta, and gamma (30-100 Hz) frequency bands. We found that depressed patients had significantly higher resting theta power in the DLPFC compared to non-depressed patients [*P* = 0.017, *z* = 2.9] **(Supplementary** Fig. 1B**, 1C)**.

### Emotional Valence and Reward-Related Oscillatory Power

We examined whether emotional valence of stimuli modulated reward-related beta and alpha power. In non-depressed patients, we found a significant main effect of emotional valence on reward-related spectral power of beta [*F* = 5.8, *d.f.* = 2, *P* 0.0096] and alpha [*F* = 7.2, *d.f.* = 2, *P* = 0.0032] oscillations in the caudate, with positive stimuli evoking a significantly higher power increase compared to negative [*P* = 0.039 beta, *P* = 0.0010 alpha] and neutral [*P* = 0.035 beta, *P* = 0.025 alpha] stimuli **(Fig. 3)**.

In depressed patients, we found a significant main effect of emotional valence on reward-related alpha power in the DLPFC [*F* = 4.1, *d.f.* = 2, *P* 0.0496], with a greater increase in alpha power seen following positive reward stimuli compared to neutral reward stimuli [*P* = 0.032]. We found no significant difference between DLPFC alpha power response to positive and negative reward stimuli [*P* = 1.9].

### Antidepressant Medication Status

We were also interested in how antidepressant medications (ADMs) may impact reward-related oscillatory power. In the DLPFC, reward-related beta power increase was significantly attenuated in patients who were taking ADMs [*P* = 3.2E-03, *z* = -3.2] **(Fig. 6B)**. We also examined the interaction between depression and antidepressant medication status and its impact on reward-related power. Since only one patient who had caudate recordings was taking ADMs while meeting the criteria for depression, we excluded caudate data from this analysis. We found a significant interaction effect between depression and ADM status on DLPFC alpha power [*F* = 23, *d.f.* = 1, *P* 1.5E-05] following reward feedback but not beta power [*F* = 4.9, *d.f.* = 1, *P* = 0.059], while depression status alone had a significant main effect on beta [*F* = 6.8, *d.f.* = 1, *P* = 0.022] power **(Fig. 6C)**.

**Figure 5.**
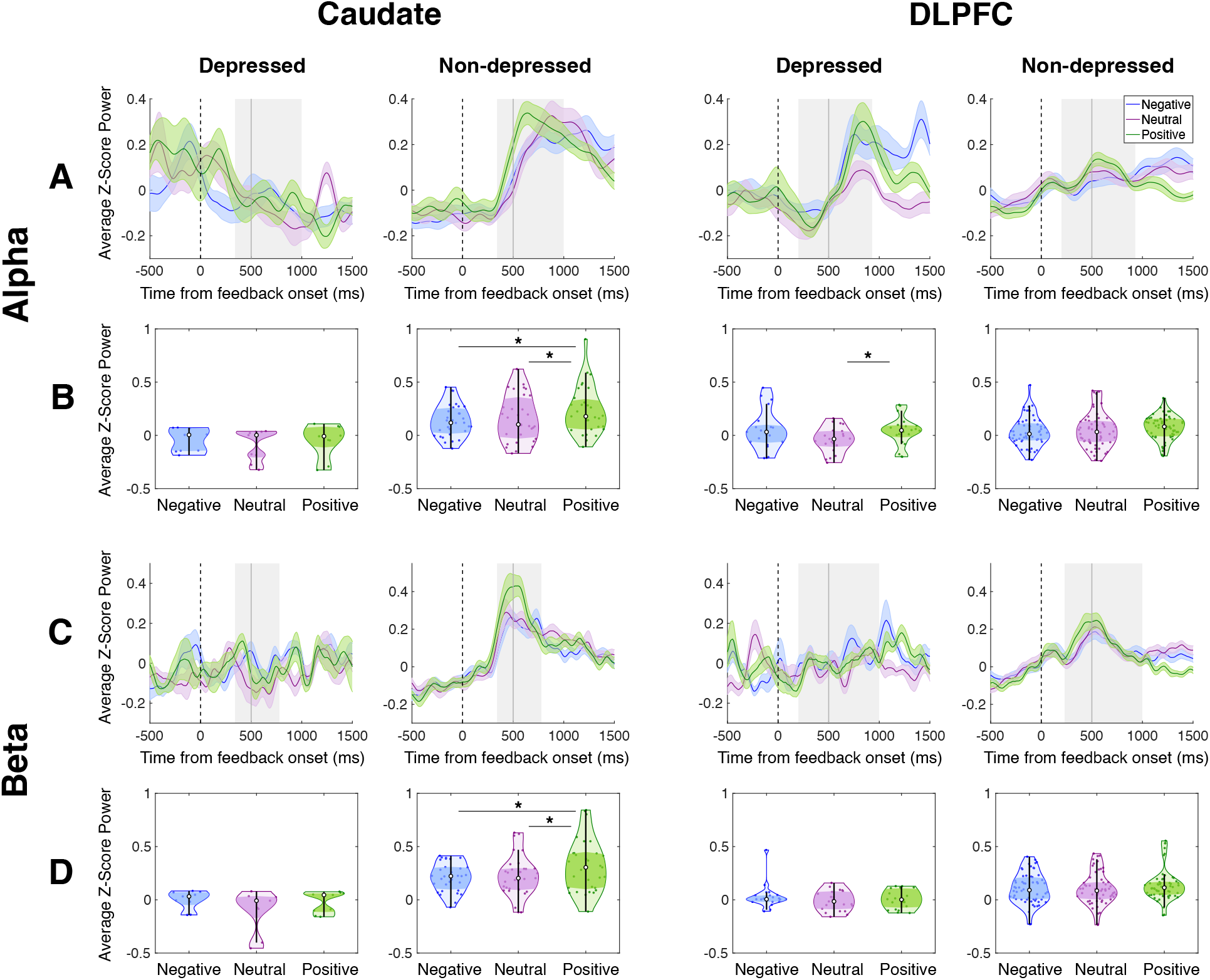
Effects of emotional valence of feedback stimuli on caudate and DLPFC alpha and beta power during reward feedback. From left to right, caudate and DLPFC data is shown in the first two columns and the last two columns respectively. Alpha and beta power data is shown in rows A-B and C-D respectively. **(A and C)** Time courses of average z-scored alpha (A) and beta (C) power during reward feedback for correct trials for depressed and non-depressed patients. Shading around the average line indicates standard error. Dotted line indicates feedback onset and solid line indicates end of feedback presentation. Gray shaded boxes indicate time clusters where band-specific power was significantly different between correct and incorrect power across all subjects. **(B and D)** Violin plots comparing average z-scored alpha (B) and beta (D) power for correct trials with negative, neutral, and positive emotional stimuli for depressed (left) and non-depressed (right) patients. White circle indicates median, middle shaded region indicates interquartile range, and black line indicates the data minimum and maximum excluding outliers. Caudate alpha and beta power following reward feedback for positive emotional stimuli was significantly higher than caudate power following reward feedback for negative and neutral emotional stimuli. DLPFC alpha power was significantly higher following reward feedback for positive emotional compared to neutral emotional stimuli * = p <0.05. DLPFC = dorsolateral prefrontal cortex.

**Figure 6.**
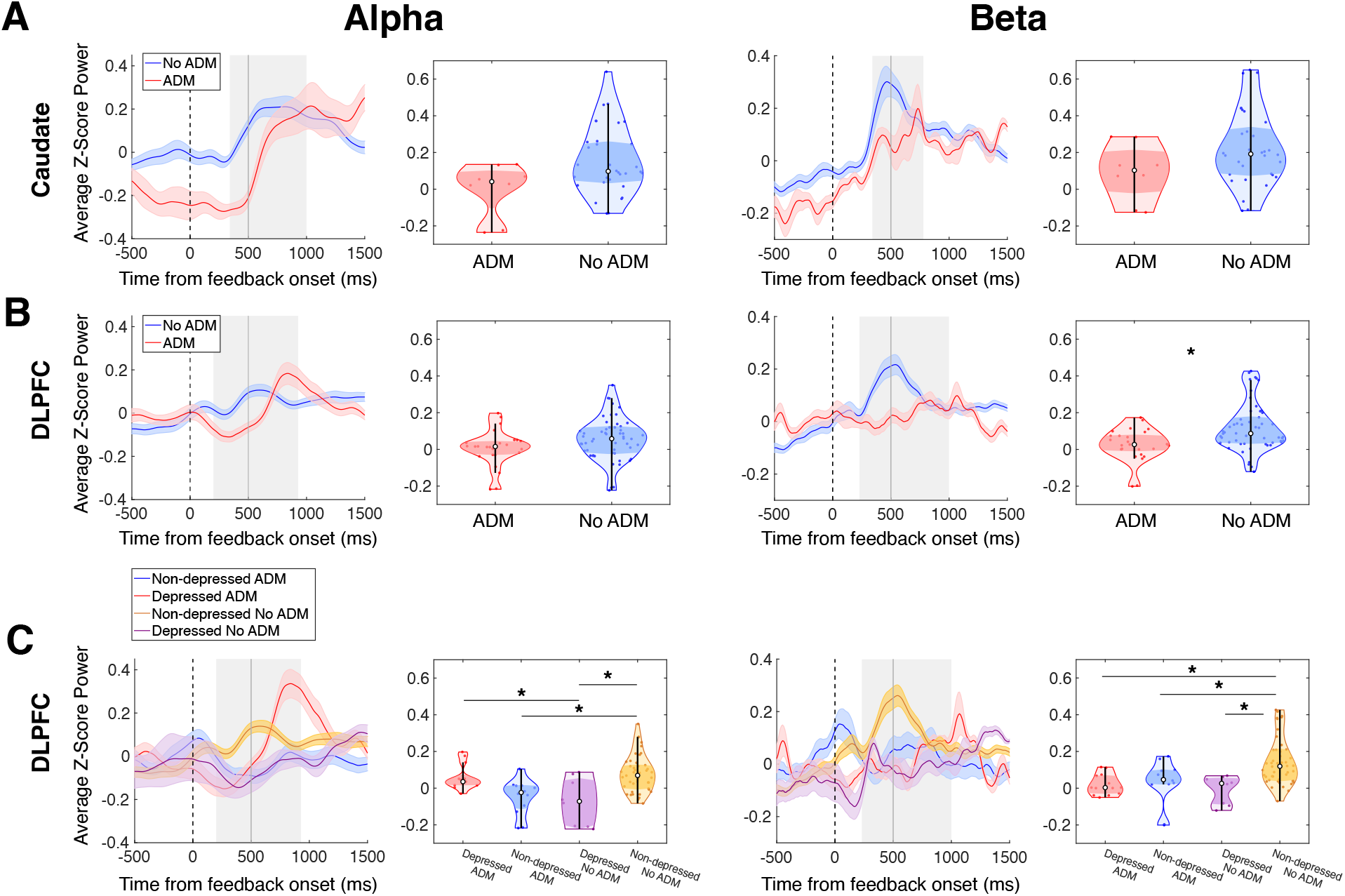
ADM status and depression alter alpha and beta power during reward feedback. From left to right, alpha and beta power data is shown in the first two columns and the last two columns respectively. **(A and B)** (Left) Time courses of average z-scored alpha and beta power in the caudate (A) and DLPFC (B) during reward feedback for correct trials, separated by ADM status. Shading around the average line indicates standard error. Dotted line indicates feedback onset and solid line indicates end of feedback presentation. Gray shaded boxes indicate time clusters where band-specific power was significantly different between correct and incorrect power across all subjects. (Right) Violin plots comparing average correct z-scored alpha and beta power in the caudate (A) and DLPFC (B) across patient groups based on ADM status. White circle indicates median, middle shaded region indicates interquartile range, and black line indicates the data minimum and maximum excluding outliers. DLPFC beta power was significantly higher for patients not taking ADMs compared to patients taking ADMs. **(C)** (Left) Time courses of average z-scored DLPFC alpha and beta power during reward feedback for correct trials, separating subjects by depression and ADM status. (Right) Violin plots comparing average z-scored DLPFC alpha and beta power across patient groups based on depression and ADM status. Depressed patients who were not taking ADMs exhibited significantly lower beta and alpha power in the DLPFC following reward feedback compared to non-depressed patients of the same ADM status. Within non-depressed patients, patients who were not taking ADMs had higher reward-related beta and alpha power compared to those who were taking ADMs. Depressed patients who were taking ADMs showed higher reward-related alpha power in the DLPFC compared to depressed patients who were not on ADMs and lower reward-related beta power compared non-depressed patients who were not taking ADMs. * = p <0.05. DLPFC = dorsolateral prefrontal cortex.

Pairwise, depressed patients who were not taking ADMs exhibited significantly lower beta [*P* = 0.0089] and alpha [*P* = 0.0013] power in the DLPFC following reward feedback compared to non-depressed patients of the same ADM status **(Fig. 6C)**. Within non-depressed patients, patients who were not taking ADMs also had higher reward-related beta [*P* = 0.032] and alpha [*P* = 0.0014] power compared to those who were taking ADMs. Depressed patients who were taking ADMs exhibited higher reward-related alpha power in the DLPFC compared to depressed patients who were not on ADMs [*P* = 0.036] and significantly lower reward-related beta power compared to non-depressed patients who were not taking ADMs [*P* = 0.0090] **(Fig. 6C)**.

We repeated this two-way ANOVA with DLPFC resting data and found a significant effect of depression on delta [*F* = 7.2, *d.f.* = 1, *P* = 0.045] and theta [*F* = 9.6, *d.f.* = 1, *P* = 0.013] band spectral power. Pairwise post-hoc comparisons showed that, between patients who were on ADMs, depressed patients exhibited significantly higher DLPFC power in the delta [*P* = 0.014] and theta [*P* = 0.025] bands than non-depressed patients. However, the interaction between depression and ADM status did not have a significant effect on DLPFC spectral power in any frequency band **(Supplementary** Fig. 1B**, 1C)**.

## Discussion

In this study, we investigated whether depression modulates spectral power signals during reward feedback in an emotional working memory task. We report that corticostriatal alpha and beta band neural oscillations are elevated during reward and that these signals are modulated by depression, ADMs, and the emotional valence of stimuli. Our primary observation was that alpha and beta power in the caudate and DLPFC increased following the onset of rewarding feedback for correct responses but not during feedback for incorrect responses. In the beta band, this reward-related increase was attenuated in patients with depression. We also found significant correlations between caudate beta power during positive feedback and total BDI-II score as well as anhedonia and somatic factor subscores. While non-depressed subjects had greater caudate alpha and beta power elevations for positive compared to neutral and negative valence stimuli, depressed subjects did not. Finally, antidepressant medications modulated DLPFC alpha power in a manner dependent on depression.

Our findings of positive feedback-related elevations in alpha and beta power are consistent with previous studies that have examined the role of frontal and corticostriatal beta band neural oscillations during cognitive tasks. A study using scalp EEG found that elevations in DLPFC beta power were associated with monetary reward in a spatial working memory task.^37^ Intracranial EEG has previously been used to demonstrate caudate and DLPFC beta power increase following reinforcement feedback for correct responses in a reinforcement learning task, and DLPFC beta power during feedback has been observed to correlate with learning.^39^ Previous studies examining the relationship between beta power and reward prediction error have suggested that an increase in beta oscillations in the DLPFC may be indicative of attentional control as well as neural processes in which the DLPFC facilitates communication with other brain regions to adapt to task requirements.^23,50^ This perspective is supported by work in non-human primates that has shown that prefrontal cortex beta oscillations are involved in the top-down regulation of the striatum during learning and attentional effort.^51–53^ Previous studies have suggested that dorsal striatum neural firing contains information about action value while DLPFC neural firing contains more information about action selection, suggesting that caudate may provide updated value information to DLPFC to guide action selection.^39,54^ The reward-related beta power elevations we observed in these structures may reflect similar processes. Importantly, we found that reward-related elevations in caudate alpha and beta power were attenuated in depressed individuals.

Activation of fronto-striatal circuits, which encompass the DLPFC and caudate nucleus, has previously been shown to play an important role in reward processes.^13,55^ Altered front-striatal activation and connectivity during reward anticipation and processing have consequently been established as prominent neural correlates of depression symptoms.^7,13^ In reward-based learning studies, depression has been associated with reduced neural activation in frontal and striatal regions during anticipation of rewards and losses^7,13^ as well as reduced functional connectivity throughout fronto-striatal circuitry during both reward anticipation and reward delivery.^13^ Subjects with depression have also been found to exhibit dysfunctional behavioral responses to reward, with reduced sensitivity to rewards and heightened maladaptive responses to punishment.^56^ Likewise, depression has also been linked to encoding biases toward negative outcomes during reinforcement learning^57^ and negative emotional stimuli.^34^

Our observations of a negative correlation between depression severity and subjects’ corticostriatal beta power response to rewarding feedback stimuli are consistent with these findings. Given its specific relationship to the experience and anticipation of reward, anhedonia is thought to be most directly linked to dysfunctional reward signaling in depression.^13,15,17,20,58^ Our finding that more severe anhedonia symptoms correlate with attenuated reward-related beta power in the caudate supports this hypothesis and provides further evidence that depression alters striatal signaling during reward.

Although changes in emotional arousal and an increased bias toward negative affect have been widely reported in patients with depression, there are conflicting findings on how depression alters brain activity during emotional processing. Previous literature has generally found that patients with depression exhibit enhanced activation of prefrontal and limbic structures in emotion-sensitive networks when presented with negative emotional stimuli.^28,30^ Some studies have shown that prefrontal structures involved in cognitive control exhibit attenuated activity during emotion processing in depressed individuals.^29,30^ Compared to controls, patients with depression have also been found to exhibit reduced activation in the caudate, amygdala, and hippocampus when responding to emotional stimuli compared to neutral stimuli, which may additionally indicate a decreased capacity for emotional arousal.^28^ Our observation that non-depressed patients exhibit a significantly greater increase in caudate alpha and beta power in response to positive word stimuli during correct trial feedback supports these previous findings in regard to the caudate’s role in emotional processing.^28^ We found that depression dampens the differences in caudate alpha and beta power response to positive, neutral, and negative word stimuli feedback, which is inconsistent with previous studies that have found changes in neural activation specific to negative emotional stimuli in patients with depression.^28,30^ Thus, our results may indicate that caudate reward signaling is enhanced by positive emotional stimuli, but this enhancement is blunted in patients with depression. On the other hand, we also found that depressed patients exhibited a greater increase in DLPFC alpha power following positive feedback compared to neutral stimuli feedback, while there was no significant difference between alpha power increase between positive and negative stimuli. This finding could suggest that when positive emotional enhancement of corticostriatal reward signaling remains intact in depressed patients, it is also accompanied by heightened sensitivity to negative stimuli.

We also found that ADM status alone generally blunted reward-related spectral power, with patients who were taking ADMs exhibiting lower beta power in the DLPFC following reward feedback for correct responses. Within non-depressed patients, ADMs lowered reward-related alpha and beta power in the DLPFC. Our analysis of the interaction between depression and ADM status, however, seems to suggest that the effects of ADMs may differ based on whether patients are exhibiting depression symptoms. In the DLPFC, ADMs restored reward-related alpha power elevations in depressed patients but attenuated them in patients who were not depressed. Similarly, DLPFC reward-related beta power in patients with depression who were not taking ADMs was significantly lower than that of non-depressed patients who were not on ADMs, but ADMs eliminated this difference in patients with depression.

There have been conflicting conclusions on how different ADMs affect reward representation: while some studies have found that short-term administration of ADMs has an adverse effect on reward processing^59^ and striatal reward circuitry activation,^60^ others have found that longer-term administration of ADMs increased activation of a widespread network of structures involved in reward and effort learning.^61^ Of note, these previous studies investigated the impact of ADMs on reward processing in healthy subjects. Our findings suggest that the impact of ADMs on reward processing may be modulated by depression.

Another prominent difference between our study and previous investigations into the relationship between depression and reward signaling is that subjects did not rely on reward feedback to inform future responses, nor did our task measure reward prediction error and neural signals associated with anticipation of reward. Our observation of a similar increase in beta power for correct trial feedback as the increase in beta power that other studies have found to occur following informative feedback stimuli, such as unexpected rewards,^50^ could suggest that corticostriatal beta power increase is inherent to reward receipt.

Limitations of this work include that all participants in our study were neurosurgical candidates, thus we were unable to compare our dataset to age-and gender-matched healthy control subjects. All subjects were diagnosed with either Parkinson’s disease or essential tremor, both of which are disorders that commonly occur with other comorbidities as well as nonmotor symptoms such as cognitive impairment. Parkinson’s disease patients are known to exhibit altered beta band neural oscillations in the cortico-striato-thalamo-cortico (CSTC) motor circuits that have been linked to their motor symptoms. Because severe depression is a contraindication to deep brain stimulation surgery for movement disorders, all depressed subjects in our study had mild to moderate depression. Further research will be necessary to examine the impact of depression severity on these findings.

Overall, this work provides evidence for the disturbance of reward-related alpha and beta band neural oscillations by depression and ADM status in a population of patients undergoing neurosurgical procedures. These findings have important implications for guiding future therapeutic developments for treatment-resistant depression (TRD). Deep brain stimulation, for instance, has generated significant interest as a treatment option for severe TRD. Our findings implicate alpha and beta power attenuation within corticostriatal circuits as a potential therapeutic target for adaptive stimulation techniques aimed at depressive symptoms such as anhedonia.

