## Supplementary Figure 1 for "Depression Attenuates Caudate and Dorsolateral Prefrontal Cortex Alpha and Beta Power Response to Reward"

### Supplementary Material

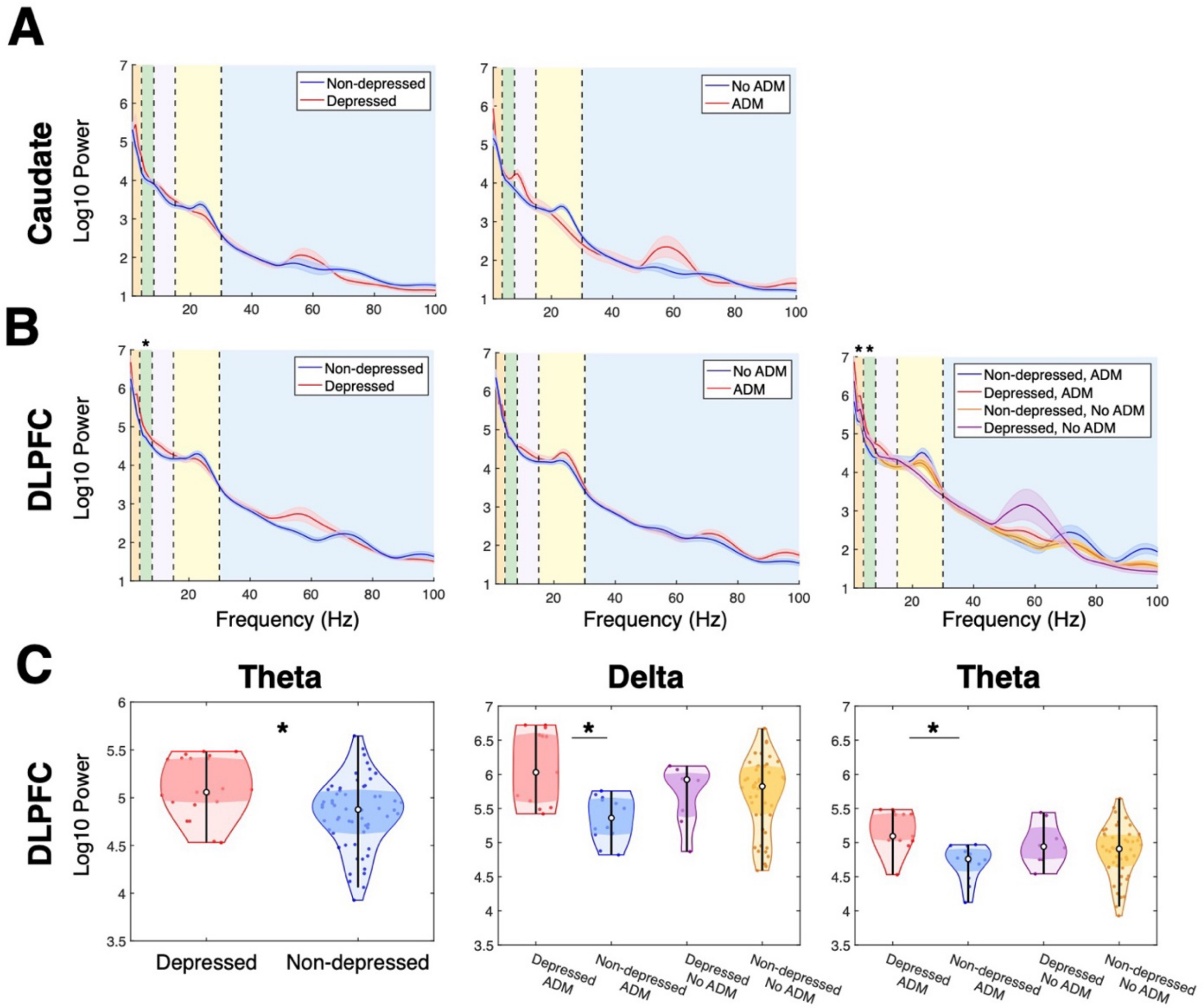


**Supplementary Figure 1. Caudate and DLPFC resting-state power by depression and ADM status. (A)** Average caudate resting power (log-transformed by base 10) separated by depressed and non-depressed patients (left) and patients who are taking versus not taking ADMs (right). **(B)** Average DLPFC resting power separated by depressed and non-depressed patients (left), by patients who are taking versus not taking ADMs (center), and by both depression and ADM status (right). **(C)** (Right) Violin plots comparing average log-transformed resting DLPFC theta power between depressed and non-depressed patients. White circle indicates median, middle shaded region indicates interquartile range, and black line indicates the data minimum and maximum excluding outliers. Depressed patients had significantly higher resting DLPFC theta power compared to non-depressed patients. (Center) Violin plots comparing average log-transformed resting DLPFC delta power across patient groups, separating subjects by depression and ADM status. (Right) Violin plots comparing average log-transformed resting DLPFC theta power across patient groups, separating subjects by depression and ADM status. Within patients who were taking ADMs, depressed patients showed significantly higher resting DLPFC delta and theta power.
